# Effects of α-tubulin acetylation on microtubule structure and stability

**DOI:** 10.1101/516591

**Authors:** Lisa Eshun-Wilson, Rui Zhang, Didier Portran, Maxence Nachury, Dan Toso, Thomas Lohr, Michele Vendruscolo, Massimiliano Bonomi, James S. Fraser, Eva Nogales

## Abstract

Acetylation of K40 in α-tubulin is the sole post-translational modification to mark the luminal surface of microtubules. It is still controversial whether its relationship with microtubule stabilization is correlative or causative. We have obtained high-resolution cryo-electron microscopy reconstructions of pure samples of αTAT1-acetylated and SIRT2-deacetylated microtubules to visualize the structural consequences of this modification and reveal its potential for influencing the larger assembly properties of microtubules. We modeled the conformational ensembles of the unmodified and acetylated states by using the experimental cryo-EM density as the structural restraint in molecular dynamics simulations. We found that acetylation alters the conformational landscape of the flexible loop that contains αK40. Modification of αK40 reduces the disorder of the loop and restricts the states that it samples. We propose that the change in conformational sampling that we describe, at a location very close to the lateral contacts site, is likely to affect microtubule stability and function.

## INTRODUCTION

Microtubules (MTs) are essential cytoskeletal polymers important for cell shape and motility and critical for cell division. They are built of αβ-tubulin heterodimers that assemble head-to-tail into ~13 polar protofilaments (PFs), which associate laterally to form a hollow tube^1^. Lateral contacts involve key residues in the so-called M-loop (between S7 and H9) in one tubulin monomer and the H2-H3 loop and β-hairpin structure in the H1’-S2 loop of the other tubulin subunit across the lateral interface. These lateral contacts are homotypic (α-α and β-β contacts), except at the MT “seam”, where the contacts are heterotypic (α-β and β-α contacts)^2^. MTs undergo dynamic instability, the stochastic switching between growing and shrinking states^3,4^. These dynamics are highly regulated *in vivo* by multiple mechanisms that affect tubulin and its interaction with a large number of regulatory factors.

One mechanism that cells can use to manipulate MT structure and function involves the post-translational modification of tubulin subunits. Through the spatial-temporal regulation of proteins by the covalent attachment of additional chemical groups, proteolytic cleavage or intein splicing, post-translational modifications (PTMs) can play important roles in controlling the stability and function of MTs^5^. Most of tubulin PTMs alter residues within the highly flexible C-terminal tail of tubulin that extends from the surface of the MT and contributes to the binding of microtubule-associated proteins (MAPs)^6,7^. These PTMs include detyrosination, Δ2-tubulin generation, polyglutamylation, and polyglycylation^8^. However, acetylation of α-tubulin on the K40 stands out as the only tubulin PTM that localizes to the inside of the MT, within a loop of residues P37 to D47, often referred to as the αK40 loop. This modification is carried out by α-tubulin acetyltransferase αTAT1 and removed by the NAD^+^-dependent deacetylase SIRT2 and by HDAC6^6,9^. How the enzymes interact with the αK40 luminal loop, and whether this “hidden PTM” has a causative or correlative effect on MT properties remain elusive.

Shortly after its discovery over 30 years ago^10^, acetylation of αK40 was found to mark stable, long-lived (t_1/2_ >2 hours) MT subpopulations, including the axonemes of cilia and flagella or the marginal bands of platelets^6,9^, and to protect MTs from mild treatments with depolymerizing drugs, such as colchicine^11^ and nocodazole^12^. Multiple studies have shown that reduced levels of αK40 acetylation cause axonal transport defects associated with Huntington’s disease, Charcot-Marie-Tooth disease, amyotrophic lateral sclerosis, and Parkinson’s disease^13–16^. These defects can be reversed by restoring αK40 acetylation levels^17^. On the other hand, elevated levels of αK40 acetylation promote cell-cell aggregation, migration and tumor reattachment in multiple aggressive, metastatic breast cancer cell lines^18,19^.

Whether acetylated MTs are stable because they are acetylated or whether stable structures are better at acquiring this modification remains a point of contention. For example, a previous study showed that acetylation did not affect tubulin polymerization kinetics *in vitro*^20^. However, this study was confounded by two factors: (i) microtubules acetylated by flagellar extract were compared to native brain tubulin, which is approximately 30% acetylated and (ii) only a single round of polymerization/depolymerization was performed after *in vitro* acetylation, which is insufficient to remove αTAT1 or other MAPs. Thus, the results of this study may be limited by the purity and preparation of the sample. Our previous structural work also found no significant differences between 30% acetylated and 90% deacetylated MTs at a resolution of ~9 Å, particularly at the modification site, the αK40 residue within the αK40 loop, which was invisible in both cases due to the intrinsic disorder and/or to the remaining heterogeneity of the loop (i.e. that study may have been limited by the low purity of the samples)^9^. More recent *in vitro* studies, using pure samples of 96% acetylated and 99% deacetylated MTs, argue that αK40 acetylation induces a structural change that improves the flexibility and resilience of MTs^21,22^. These studies find that acetylated MTs maintain their flexural rigidity, or persistence length, after repeated rounds of mechanical stress, while deacetylated MTs show a 50% decrease in rigidity and are 26% more likely to suffer from complete breakage events^21,22^.

Since the αK40 residue is less than 15 Å away from the lateral interface between protofilaments, a possible model for the molecular mechanism of acetylation is that it alters inter-protofilament interactions by promoting a conformation of the αK40 loop that confers flexural rigidity, thus increases its resistance to mechanical stress—a phenomenon called protofilament sliding^21,22^. Molecular dynamics simulations have suggested a model where αK40 forms a stabilizing salt bridge with αE55 within the core of the α-tubulin monomer that in turn stabilizes αH283 within the M-loop of its neighboring α-tubulin monomer^23^. Another study proposed that αK40 acetylation may specify 15-PF MTs, which are known to be 35% stiffer than 13PF MTs and more effective at forming microtubule bundles^24^.

Given the uncertainties remaining concerning the effect of αK40 acetylation on MTs, we decided to characterize the conformational properties of the αK40 loop in the acetylated and deacetylated MTs that could have an effect on MT structure and properties. To that end, we produced near atomic-resolution cryo-EM maps of 96% acetylated (Ac^96^) and 99% deacetylated (Ac^0^) MTs. By improving sample purity, we were able to visualize more density for the αK40 loop in the acetylated state. Using new molecular dynamics methods, we found that acetylation shifts the conformational landscape of the αK40 loop by restricting the range of motion of the loop. In contrast, in the Ac^0^ state, the αK40 loop extends deeper into the lumen of the MT, and samples a greater number of conformations. These motions are likely to increase the accessibility of the loop to αTAT1, in agreement with the hypothesis that αTAT1 acts by accessing the MT lumen^25^, and likely influence lateral contacts, in agreement with the causative effect of acetylation on the mechanical properties of microtubules^21^.

## RESULTS AND DISCUSSION

### High-resolution cryoEM reconstructions of pure acetylated (Ac^96^) and deacetylated (Ac^0^) MTs

Using recent biochemical schemes designed to enrich for specific acetylation states^21^, we generated Ac^96^ and Ac^0^ MTs for use in our cryo-EM studies. We prepared cryo-EM samples as previously described^2,26^ of Ac^96^ and Ac^0^ MTs in the presence of end-binding protein 3 (EB3). EB3 served as a fiducial marker of the dimer that facilitated alignment of MT segments during image processing^27^. Table S1 summarizes the data collection, refinement, and validation statistics for each high-resolution map we visualized (see also Supplemental Figures 1 and 2). Using the symmetrized MT reconstruction, which takes advantage of the pseudo-helical symmetry present in the MT, we extracted a 4×3 array of dimers for further B-factor sharpening, refinement^28^, and model-building (Figure 1a). This array includes all possible lateral and longitudinal non-seam contacts for the central dimer, which was later extracted for model building and map analysis (Figure 1b,1c).

**Figure 1.**
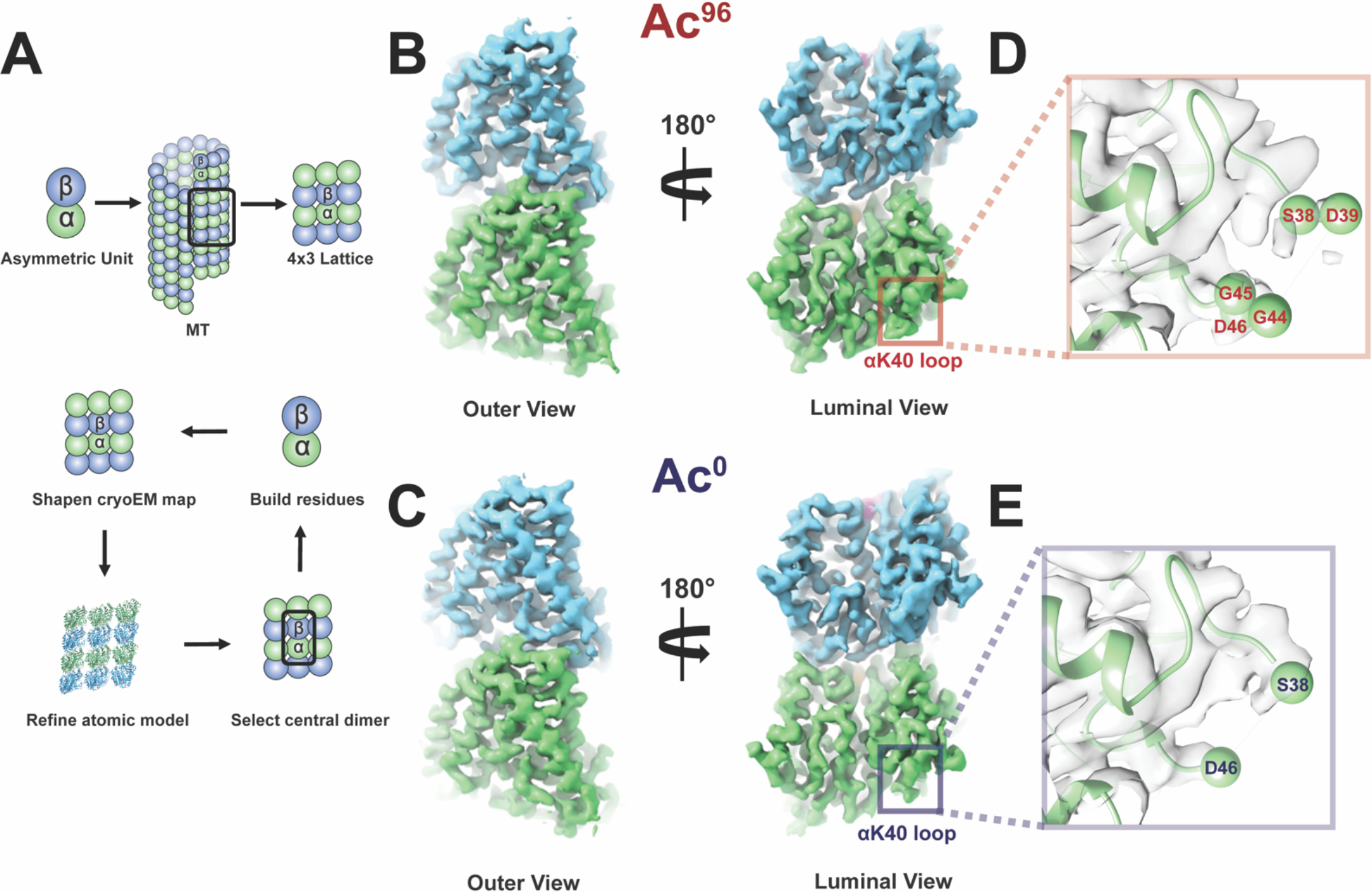
High-resolution maps of 96% acetylated (Ac^96^) and <1% acetylated (Ac^0^) microtubules. **(a)** Schematic of the model-building and refinement process in PHENIX. We sharpened a representative 4×3 lattice, refined the corresponding atomic structure (3JAR) into our map, and extracted out the central dimer to build additional residues into the αK40 loop. We performed this process iteratively for both the Ac^96^ and Ac^0^. The structure of the Ac^96^ **(b)** and Ac^0^ **(c)** αβ-tubulin heterodimers, respectively, are shown from the outer and luminal views with close-ups of αK40 loop in each state **(d)** and **(e)** low-pass filtered to 3.7 Å.

The αK40 loop has been poorly resolved in previous EM reconstructions, and existing models contain a gap between residues Pro37 and Asp48 (Supplemental Figure 3a)^25^. While the loop has been resolved in a number of X-ray crystallographic structures, the conformations stabilized in the crystal lattice are likely artifacts due to the presence of calcium and/or crystal contacts (Supplemental Figure 3b). For our symmetrized maps, we were able to build residues S38-D39 and G44-D47 into the Ac^96^ state and S38 and D46-D47 into the Ac^0^ state (Figure 1d, 1e). Qualitatively, the maps suggest that the αK40 loop is slightly more ordered in the Ac^96^ state, with the protrusion of density following Pro37 extending away from or toward Asp48 in the Ac^96^ or Ac^0^ states, respectively. However, it is likely that multiple conformations of the loop, perhaps as a function of each loop’s individual position around a helical turn, are averaged together and result in the low signal-to-noise levels we observe in the map.

### Conformational differences across MT states are confirmed by non-symmetrized reconstructions

We considered the possibility that the symmetrizing procedure used to improve signal and resolution in our image analysis was averaging different αK40 loop conformation within different PFs and thus interfering with our interpretation of the loop structure in the two states. To test the hypothesis, we analyzed the non-symmetrized maps calculated with C1 symmetry for the Ac^96^ and Ac^0^ states. We extracted a full turn of 13 adjacent dimers. This full-turn map revealed additional density extending out further along the loop in the Ac^96^ state when compared to the symmetrized maps filtered to the same resolution (4 Å) (Figure 2). Furthermore, the density for the loop observed at the seam was distinct from that at the non-seam contacts. To maximize the interpretability of the subunits making non-seam contacts, we used non-crystallographic symmetry (NCS) averaging as an alternative method to increase the signal-to-noise levels in the maps. This procedure improved the density for non-Glycine backbone atoms in the αK40 loop in the Ac^96^ state, allowing us to trace an initial Cα backbone for this region, while in the Ac^0^ state the loop remained unmodelable (Figure 2c, 2d). This interpretation agrees with the qualitative difference in the density, which indicate less disorder for the Ac^96^ state than Ac^0^ state, of the traditionally symmetrized and C1 maps.

**Figure 2.**
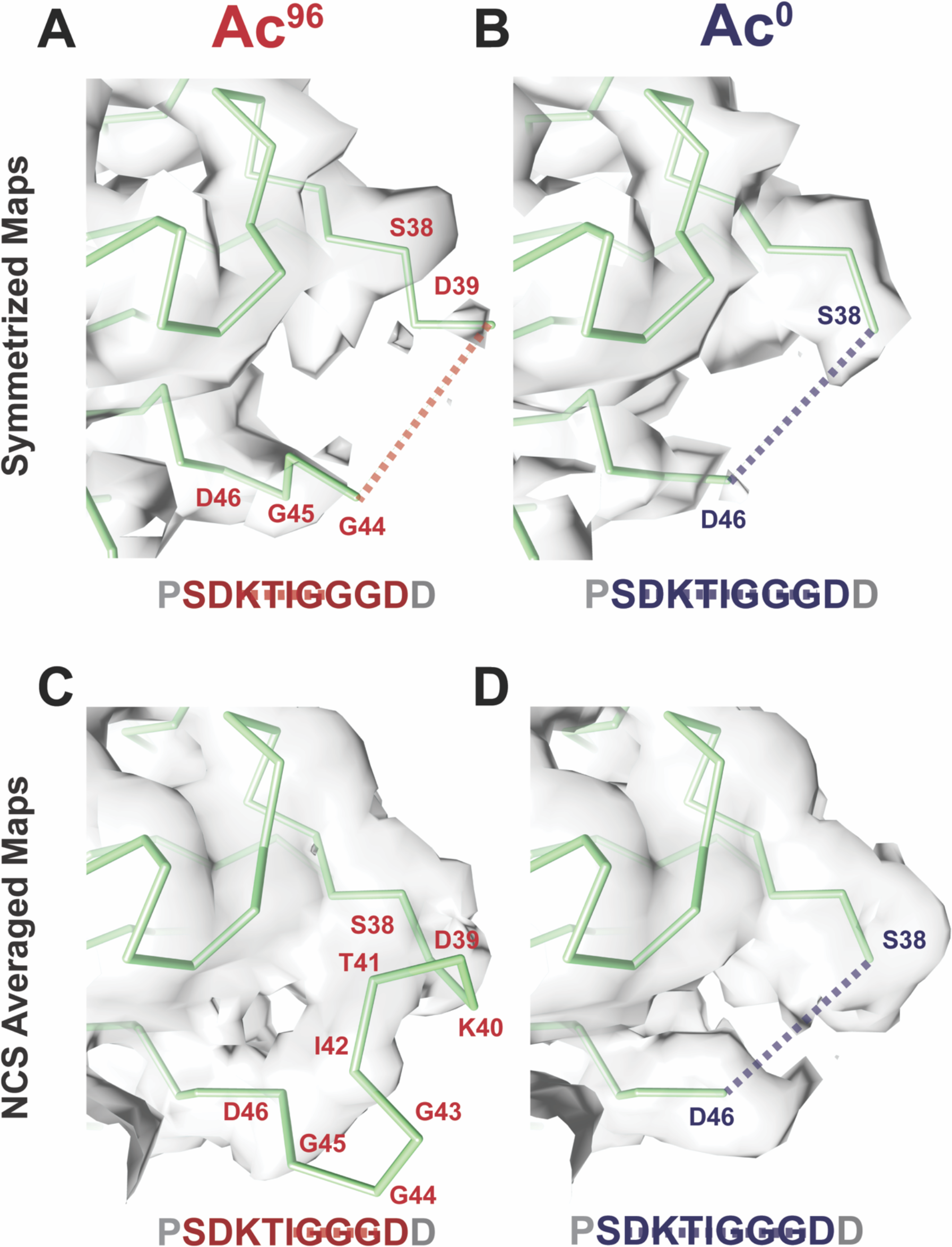
Symmetrized and NCS-averaged C1 maps of Ac^96^ and Ac^0^ microtubules reveal the αK40 loop is more ordered in the Ac^96^ state. Close-up views of the αK40 loop (P37-D47) in the **(a)** Ac^96^ and **(b)** Ac^0^ states in the symmetrized maps low-pass filtered to 4 Å and the **(c)** Ac^96^ and **(d)** Ac^0^ states in the NCS averaged C1 maps low-pass filtered to 4 Å. Dotted lines indicate missing residues.

This NCS averaging method had multiple advantages over the traditional averaging technique for pseudo-helical processing implemented in FREALIGN^26^. First, the model coordinates used for the averaging are based on the matrix of α-tubulin monomers along a full turn rather than the single α-tubulin monomer. Second, in the FREALIGN averaging approach the signal from the dimers at the seam are down-weighted, whereas NCS averaging allows us to separate the signal from the seam, and thus to deconvolute the signal from the non-seam locations. Third, this procedure also acts to low-pass filter the map to 4 Å (the high-resolution limit of the C1 map, Supplemental Figure 5), which should suppress noise from the more disordered parts of the map, including alternative conformations of the αK40 loop. Using this NCS-based approach, we were able to resolve density and build a model for three additional residues, the acetylated K40, T41, and I42. These residues pack towards the globular domain of α tubulin, consistent with the favorability of burying these relatively hydrophobic residues in the Ac^96^ state. Despite observing only very weak density, we have modeled the glycine-rich region that extends into the lumen as a tight turn, which we note is only possible due to the expanded Ramachandran space accessible to glycine residues (Figure 2c). In contrast, and despite better global resolution, we did not observe any density consistent with a stable conformation of the loop in the Ac^0^ map. Based on this result, which is consistent across the NCS-averaged and traditionally symmetrized maps, we did not build any additional residues into the Ac^0^ density (Figure 2d).

### Ensemble modeling of the loop in each state using density-restrained molecular dynamics

For regions that exhibit a high degree of disorder, like the αK40 loop, a single, static structure is a poor description of the native state. Ensemble models can help to elucidate how populations of conformations change upon perturbations, such as post-translational modifications^29,30^. To derive an ensemble of conformations representing the Ac^96^ and Ac^0^ states, we used the atomic structure built into the Ac^96^ map as the starting model to initiate metainference-based molecular dynamics (MD) simulations, which augment a standard forcefield with a term representing the density derived from the EM map^31^. In contrast to Molecular Dynamics and Flexible Fitting (MDFF) and other refinement methods that seek to converge on a single structure^32^, this method models a structural ensemble by maximizing the collective agreement between simulated and experimental maps, and accounts for noise using a Bayesian approach^33^. Initiating simulations for both the Ac^96^ and Ac^0^ states from starting models that differ only in the acetyl group and distinct input experimental density maps allowed us to test whether acetylation restricts the motion of the loop, trapping it in a tighter ensemble of conformations.

To analyze the conformational dynamics of the loop, we analyzed the root mean square fluctuations of residues 36-48 within replicas for each simulation. This analysis shows that the αK40 loop fluctuations are more restricted in the Ac^96^ state than in the Ac^0^ state (Figure 3a). Next, we analyzed the distribution of conformations adopted by the loop by analyzing the distance between K40 and the globular domain of α-tubulin (represented by L26) and by clustering together the snapshots from all replicas of both simulations based on the root mean square deviations of residues 36-48. Similar to the starting reference model, where the distance is 10.6 Å, Ac^96^ is enriched in conformations that pack close to the globular domain of the α-tubulin core (Figure 3b). These conformations, exemplified by clusters 1, 4, and 6, position the acetylated lysine to interact with residues along H1. In contrast, the Ac^0^ state favors conformations that extend towards the MT lumen, as exemplified by clusters 0, 2, 5, 7, and 8 (Figure 3b). Clusters 3, 9, 10, 11, labeled in grey, had equal numbers of frames enriched in Ac^96^ and Ac^0^ and sampled rare (<5%) extreme states on both the exposed and packed ends of the conformational spectrum (Figure 3b).

**Figure 3.**
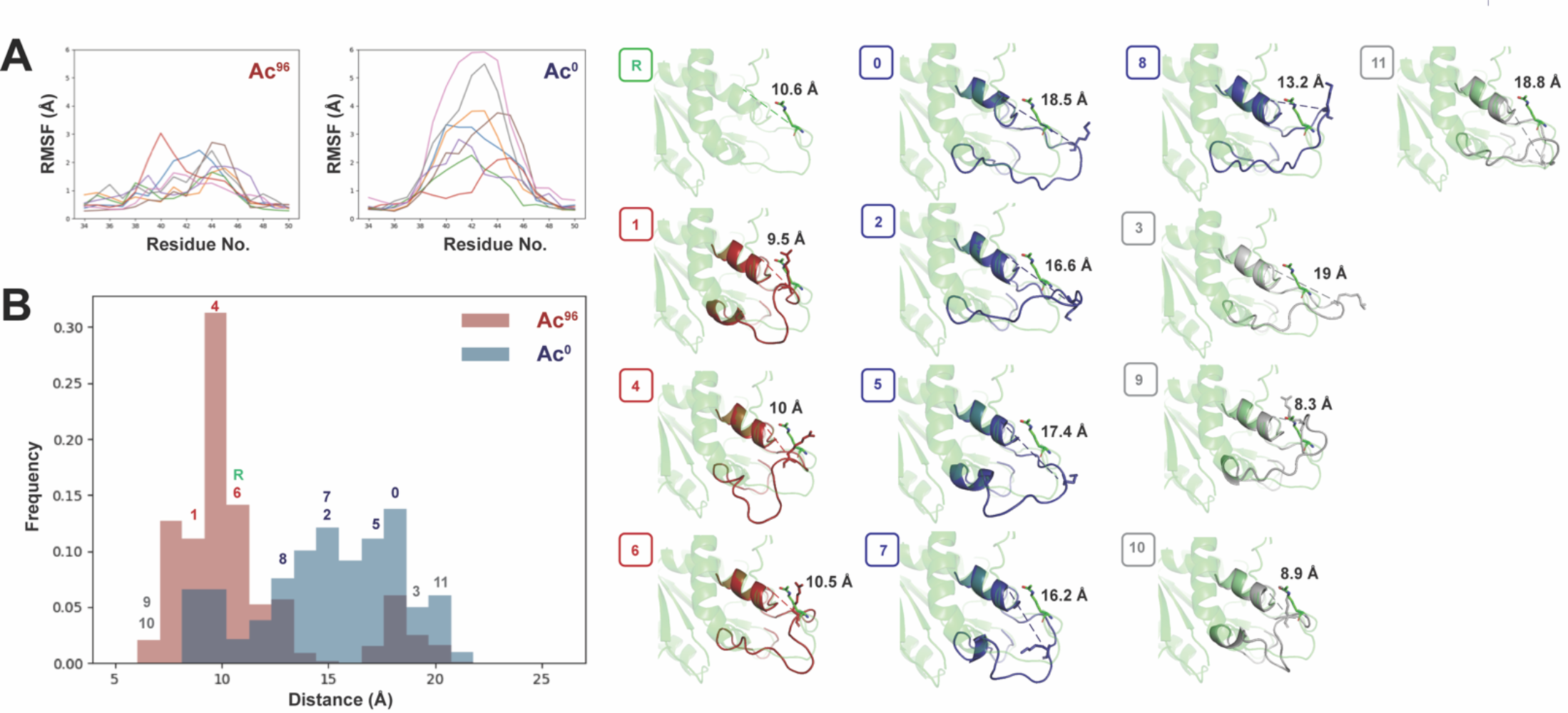
Acetylation restricts the motion and alters the conformational ensemble of the αK40 loop. **(a)** Per-residue root mean square fluctuations (RMSF) analyses were determined over the course of 12 ns for residues 34-50 the C1 maps using GROMACs in PLUMED and graphed using the MDAnalysis. (b) Ensemble modeling of the loop across Ac^96^ and Ac^0^ states using density restrained MD. Frames were classified into one of 11 clusters by conformation. Clusters either had a greater number of Ac^96^ frames (red), Ac^0^ frames (blue), or an equal number of frames from both states (grey). The reference is shown in green. The unique conformations of each of the 11 clusters are shown to the right.

These computational results are consistent with the visual analysis of the density for both the NCS and traditionally symmetrized maps, which indicated that the loop is more ordered after acetylation. The residual disorder identified by the simulations using the Ac^96^ map may be important for de-acetylation by SIRT2. On the other hand, the increased flexibility we observe for the Ac^0^ state suggests a potential mechanism by which αTAT1 could acetylate K40. Previous proposals argue that acetylation can occur from the outside or inside of the lumen^25^. However, to catalyze the modification, a flexible region within αTAT1 would have to extend approximately 25 Å through a MT wall fenestration between four tubulin dimers to reach αK40, or the MT would have to undergo a major structural rearrangement in the lattice to allow αTAT1 to enter the lumen. Previous work demonstrated that the αTAT1 active site and its MT recognition surface is concave and could not stretch through the lumen^25^. Our findings support the idea that αTAT1 modifies the loop from within the lumen of the MT because the deacetylated loop samples extended structures that would be accessible to αTAT1 and because the structural rearrangement caused by acetylation is small and local to the αK40 loop.

### Acetylation induces a local structural rearrangement of the αK40 loop that promotes stability by weakening lateral contacts

Collectively our structural and MD results show that acetylation restricts the motion of the αK40 loop. These results led us to hypothesize that the change in the structural ensemble of the αK40 loop upon acetylation, while subtle and local, may affect lateral contacts. These local changes may disrupt the small lateral interface between α-tubulin subunits. The origin of this effect may be highly distributed, as we do not visualize any stable interactions between the Ac^0^ state of the loop and the globular domain. However, upon acetylation, the structural ensemble becomes more restricted and the potential for the loop to strengthen any of these interactions between monomers is lost. For example, in many of the extended conformations favored by the Ac^0^ state, K40 in a α1-monomer is close to the M-loop of the neighboring α2-monomer and may buttress the H1’-S2 loop, providing support for the vital _α1_K60:_α2_H283 lateral interaction (Figure 4). In contrast, when K40 is acetylated it packs ~10 Å closer to the globular domain of the α1-monomer, reducing the potential for inter-monomer interactions (Figure 4).

**Figure 4.**
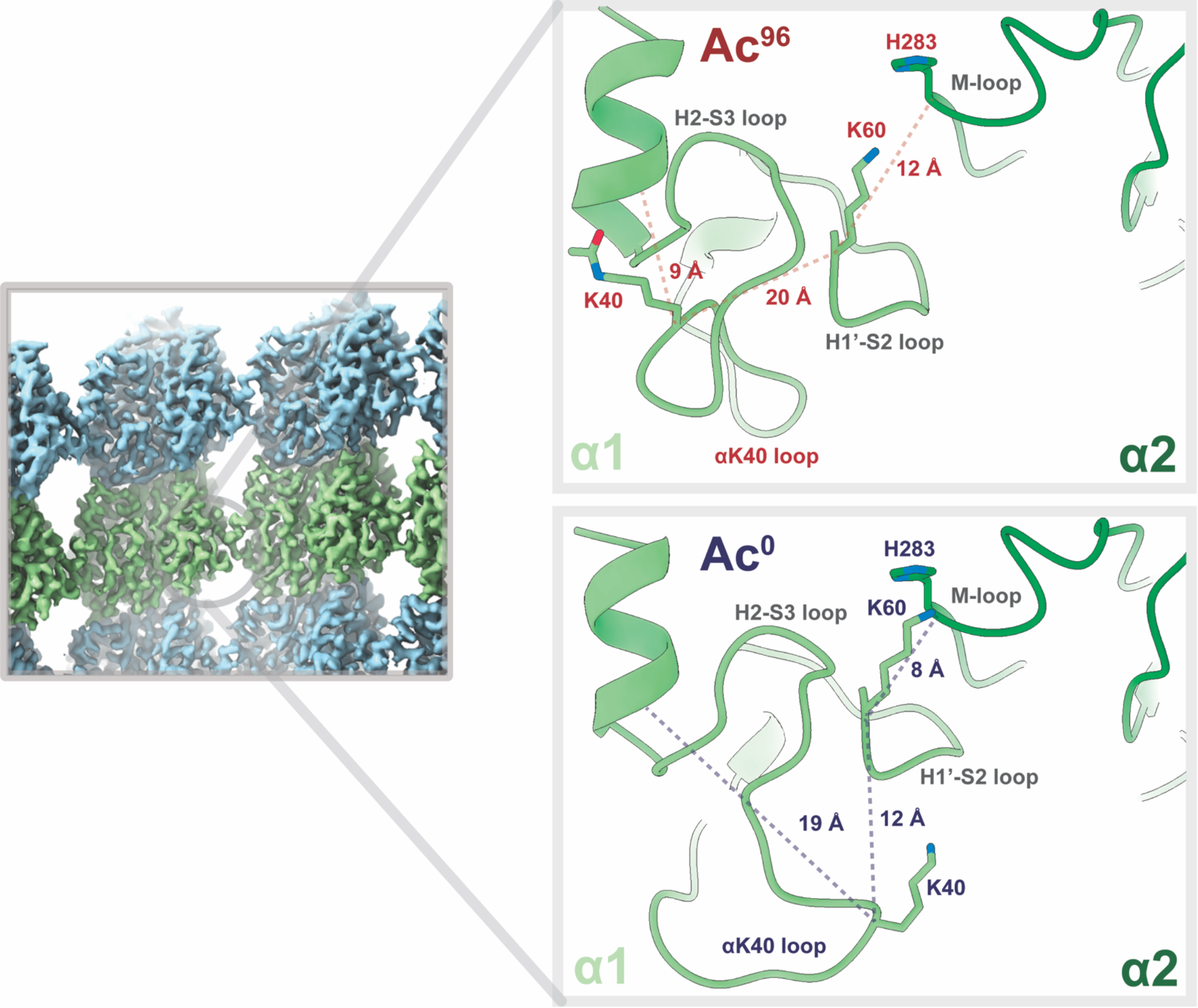
Acetylation may weaken lateral interactions. Close-up view of the lateral contacts between two α-tubulin monomers at a non-seam location (α1, light green; α2, dark green). K40 in α1 of the Ac^0^ state is 8 Å closer to the M-loop of α2 and appears to buttress the H1’-S2 loop, providing support for the vital α1K60-α2H283 lateral interaction. By contrast, that support is lost in the Ac^96^ state because the acetylated K40 now packs much closer to the hydrophobic, inner core.

We tested whether the loss of the positive charge of the lysine upon acetylation alters the electrostatic interaction energy and the hydrogen-bonding network at the lateral interface using MD simulations based on the Debye-Hückel (DH) formula^34,35^. We found that acetylation does indeed weaken lateral interactions (Supplemental Figure 4). Additionally, the Ac^0^ ensemble contains conformations with strong DH interaction energies that do not exist in the Ac^96^ ensemble (Supplemental Figure 4). While the effects of acetylation are subtle, the local effects at the lateral contacts site may have an additive effect that stabilizes the MT lattice. This idea is consistent with previous work that argues that the weakening of lateral interactions is a protective mechanism to prevent pre-existing lattice defects from spreading into large areas of damage under repeated stress—a mechanism that could be exploited by cancer cells^21,22^.

In conclusion, this comprehensive approach combines the structural insight of cryoEM with the sampling efficiency and global scope of MD to investigate how PTMs can transform a conformational ensemble^36,37^. Our high-resolution maps serve as a blueprint for the scale of conformational change and relevant degrees-of-freedom that the αK40 loop can sample with all-atomistic metainference MD^36^. We show that αTAT1 induces a site-specific electrostatic perturbation that restricts the motion of the loop. αK40 acetylation may function as an evolutionarily conserved ‘electrostatic switch’ to regulate MT stability^36,37^.

## MATERIALS AND METHODS

### Sample Preparation for Cryo-Electron Microscopy

Porcine brain tubulin was purified as previously described^38^ and reconstituted to 10 mg/ml in BRB80 buffer (80 mM 1,4-piperazinediethanesulfonic acid [PIPES], pH 6.9, 1 mM ethylene glycol tetraacetic acid [EGTA], 1 mM MgCl_2_) with 10% (vol/vol) glycerol, 1 mM GTP, and 1 mM DTT, and flash frozen in 10 μl aliquots until needed. The acetylated and deacetylated MTs (15 µM) were co-polymerized with end-binding protein 3 (EB3, 25 µM), at 37°C for ~15 min in the presence of 10% NP-40, 1mM dithiothreitol (DTT), and BRB80 buffer. The EB3 decorated MTs were added to glow-discharged C-flat holey carbon grids (CF-1.2/1.3-4C, 400 mesh, Copper; Protochips, Morrisville, NC) inside a Vitrobot (FEI, Hillsboro, OR) set at 37°C and 85% humidity before plunge-freezing in ethane slush and liquid nitrogen, respectively, as previously described^2^.

### Cryo-Electron Microscopy

Micrographs were collected using a Titan Krios microscope (Thermo Fisher Scientific, Inc., Waltham, MA) operated at an accelerating voltage of 300 kV. All cryo-EM images were recorded on a K2 Summit direct electron detector (Gatan, Pleasanton, CA), at a nominal magnification of x22,500, corresponding to a calibrated pixel size of 1.07 Å. The camera was operated in super-resolution mode, with a dose rate of ~2 e^-^ per pixel per s on the detector. We used a total exposure time of 4 s, corresponding to a total dose of 25 electrons/Å^2^ on the specimen. The data were collected semi-automatically using the SerialEM software suite^39^.

### Image Processing

Stacks of dose-fractionated image frames were aligned using the UCSF MotionCor2 software^40^. MT segments were manually selected from the drift-corrected images (acetylated dataset: 205 images, deacetylated MT dataset: 476 images) using the APPION image processing suite^41^. We estimated the CTF using CTFFIND4^42^ and converted the segments to 90% overlapping boxes (512 × 512 pixels) for particle extraction. The remaining non-overlapping region is set to 80 Å and corresponds to the tubulin dimer repeat (asymmetric unit). Consequently, there are ~13 unique tubulin dimers per MT particle. To determine the initial global alignment parameters and PF number for each MT particle, raw particles were compared to 2D projections of low-passed filtered MT models (~20 Å, 4° coarse angular step size) with 12, 13, 14 and 15 PFs^43^ using the multi-reference alignment (MSA) feature of EMAN1^44^. Finally, 13-PF MT particles (acetylated dataset: 20,256 particles, deacetylated MT dataset: 29,396) were refined in FREALIGN v. 9.11^45,46^ using pseudo-helical symmetry to account for the presence of the seam. To verify the location of the seam, we used the 40 Å shift approach to categorize MTs based on their azimuthal angle, as previously described^27^.

### Atomic Model Building and Coordinate Refinement

COOT^47^ was used to build the missing polypeptides of the αK40 loop in α-tubulin, using the available PDB 3JAR as a starting model. Successively, all novel atomic models were iteratively refined with phenix.real_space_refine into EM maps sharpened with phenix.autosharpen^28,48^. For visual comparisons between states, potential density thresholds were interactively adjusted in Coot to maximize iso-contour similarity around backbone atoms distant from the αK40 loop. For Figures 1 & 2, all densities are represented in Chimera at a threshold of 1.1.

### Molecular Dynamics Simulations

Code for map preparation, simulation execution, and analysis is available at: https://github.com/fraser-lab/plumed_em_md To prepare the cryoEM maps, we fitted the maps with a Gaussian Mixture Model (GMM) by applying a divide-and-conquer approach^33^, using generate_gmm.py and convert_GMM2PLUMED.sh. Cross-correlations to the experimental maps were greater than 0.99. All simulations were performed with GROMACS 2016.5^34^ and the PLUMED-ISDB module^49^ of the PLUMED library^50^ using the Charmm36-jul2017 forcefield^51^ with patches for acetylated lysine (aly)^52^ and the TIP3P water model. For the deacetylated simulations, the same starting model was used with a manual edit of the PDB to eliminate the acetylation (with all hydrogens replaced by GROMACS during model preparation). The initial model was minimized then equilibrated for 2ns, using prep_plumed.py. MD simulations were performed on a metainference ensemble of 8 replicas for an aggregate simulation time of 96ns for each acetylation state, using prep_plumed2.py and prep_plumed3.py. Contributions of negative scatterers (atoms OD1 and OD2 of Asp residues; OE1 and OE2 of Glu) were excluded from contributing to the predicted maps during the simulation. This modification effectively eliminates the contribution of these side chains to the agreement between density maps, in keeping with the non-existent density of negatively charged side chains in EM maps, while allowing them to contribute to the simulation through the energy function. Clustering and convergence analyses^31^ were performed and analyzed using MDAnalysis^53^.

Changes in the electrostatic interaction energies at the lateral contacts were determined using the using the Debye-Hückel (DH) formula:

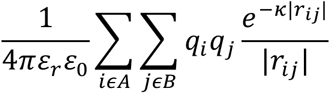

where *ϵ*_0_ is the vacuum’s dielectric constant, *ϵ*_*r*_ the dielectric constant of the solvent, *q*_*i*_ and *q*_*j*_ the charges of the *i*-th and *j*-th atoms, respectively, |*r*_*ij*_| the distance between these two atoms, and 6 is the DH parameter^35^ defined in terms of the temperature *T* and the ionic strength of the solution *I*_*s*_.

The DH energy is calculated between the following two groups of atoms, denoted as A and B in the formula above: (i) all atoms in residue range 30-60 of chain A (α1 subunit) and (ii) all atoms in residue range 200-380 of chain E (α2 subunit) in PDBs **<u>XXYA</u>** and **<u>XXYB</u>**. Residues not included in this range do not significantly contribute to the DH interaction energy between adjacent α-subunits. Parameters used in the calculation of the DH energy are: temperature (T=300K), dielectric constant of solvent (*ϵ*_*r*_=80; water at room temperature), and ionic strength (*I*_*s*_=1M).

## ACCESSION NUMBERS

All electron density maps have been deposited in the EMDB accession numbers <u>**EMD-X1**, **EMD-X2**, **EMD-X3**</u>, and <u>**EMD-X4**</u>. Atomic models are deposited in the PDB accession numbers **<u>XXYA</u>**, **<u>XXYB, XXYC</u>**, and **<u>XXYD</u>**, **<u>XXYE</u>**.

## AUTHOR CONTRIBUTIONS

D.P. performed the tubulin purification and modification to generate the Ac^0^ and Ac^96^ samples. L.E., R.Z., and D.T. performed the cryo-EM sample preparation, electron microscopy and data processing. L.E. and J.S.F. engineered the NCS-averaging technique, performed iterative model-building/refinement. L.E., J.S.F., T. L., M. V. and M.B. executed the metainference MD simulations. All authors contributed to structure interpretation, model development and manuscript writing.

## ACKNOWLEDGEMENTS

We thank P. Grob and J. Fang for cryo-EM data collection support, A. Chintangal and P. Tobias for computational support, and E. Kellogg, B. LaFrance, S. Howes, T.H.D. Nguyen, S. Pöpsel, B. Greber, and K. Morris for helpful discussions. We also acknowledge the Berkeley Bay Area Cryo-EM (BACEM) Facility and additional scientific resources at UC Berkeley. J.S.F was funded by the UCSF-UCB Sackler Faculty Exchange Program and NIGMS grant R01-GM123159. This work was funded through NIGMS grants R01-GM63072 to E.N. and the NSF grant 2016222703 and the NAS NRC Ford Foundation grant to L.E. E.N. is a Howard Hughes medical investigator.

---

**Supplemental Figure 1.**
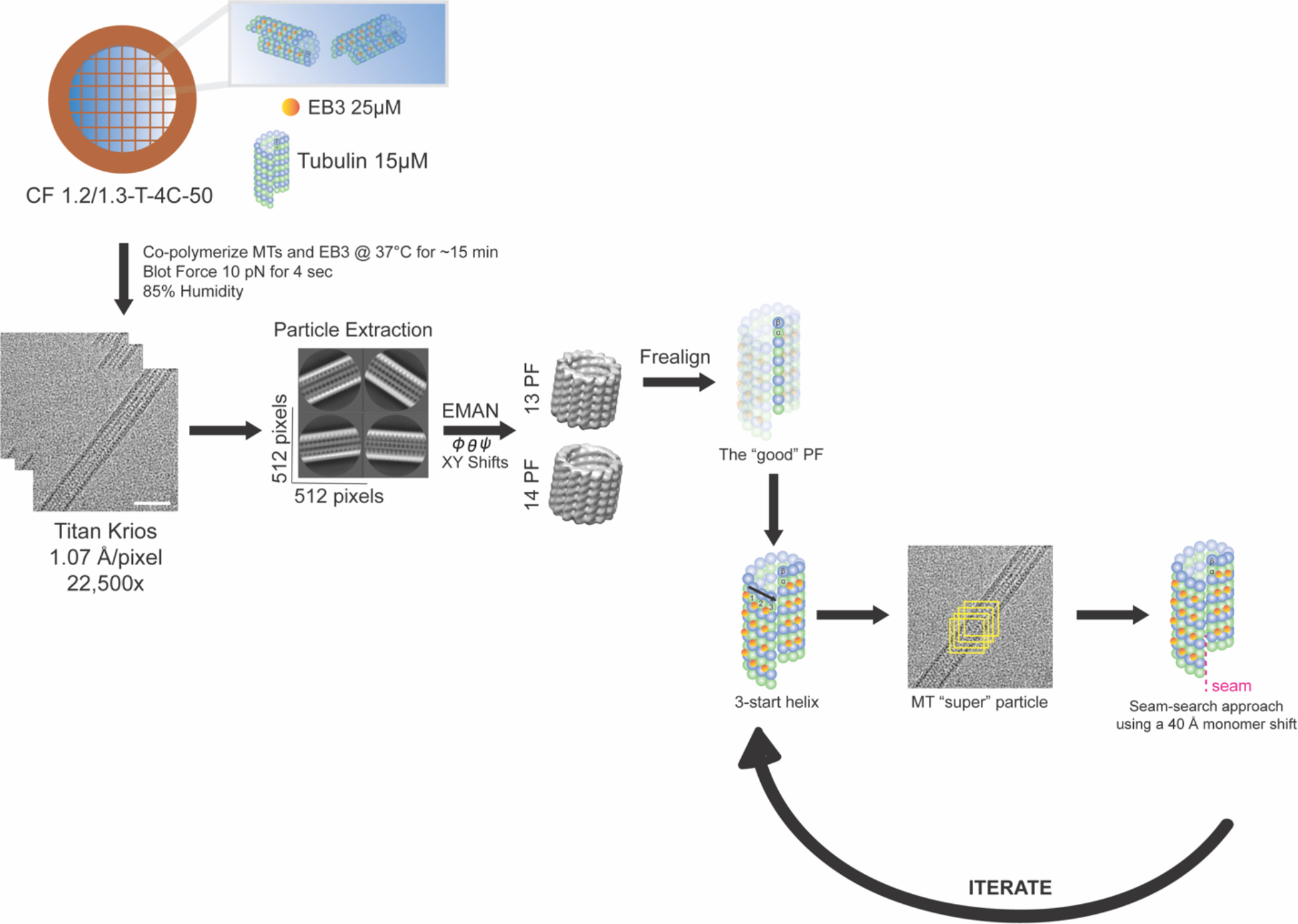
Schematic of the experimental workflow for sample preparation and pseudo-helical image processing. EB3 decorated MTs were added to glow-discharged C-flat holey carbon grids (CF-1.2/1.3-4C, 400 mesh, Copper; Protochips, Morrisville, NC) inside a Vitrobot (FEI, Hillsboro, OR) set at 37°C and 85% humidity before plunge-freezing in ethane slush and liquid nitrogen. Images were collected with the Titan Krios electron microscope (Thermo Fisher Scientific, Inc., Waltham, MA) operated at 300kV and equipped with a K2 direct detector (Gatan, Pleasanton, CA). The micrographs were collected at a nominal magnification of ×22,500. Stacks of dose-fractionated image frames were aligned using the UCSF MotionCor2 software and CTF-corrected with CTFFIND4. MT segments were manually selected and converted to 90% overlapping boxes (512 × 512 pixels) for particle extraction. The remaining non-overlapping region is set to 80 Å and corresponds to the tubulin dimer repeat (asymmetric unit). These raw particles were compared to 2D projections of low-passed filtered MT models (~20 Å, 4° coarse angular step size) with 13 and 14 PFs using the multi-reference alignment (MRA) feature of EMAN1. Next, 13-PF MT particles were refined in FREALIGN v. 9.11 using pseudo-helical symmetry to account for the presence of the seam. To verify the location of the seam, MTs were categorized based on their azimuthal angle and refined again.

**Supplemental Figure 2.**
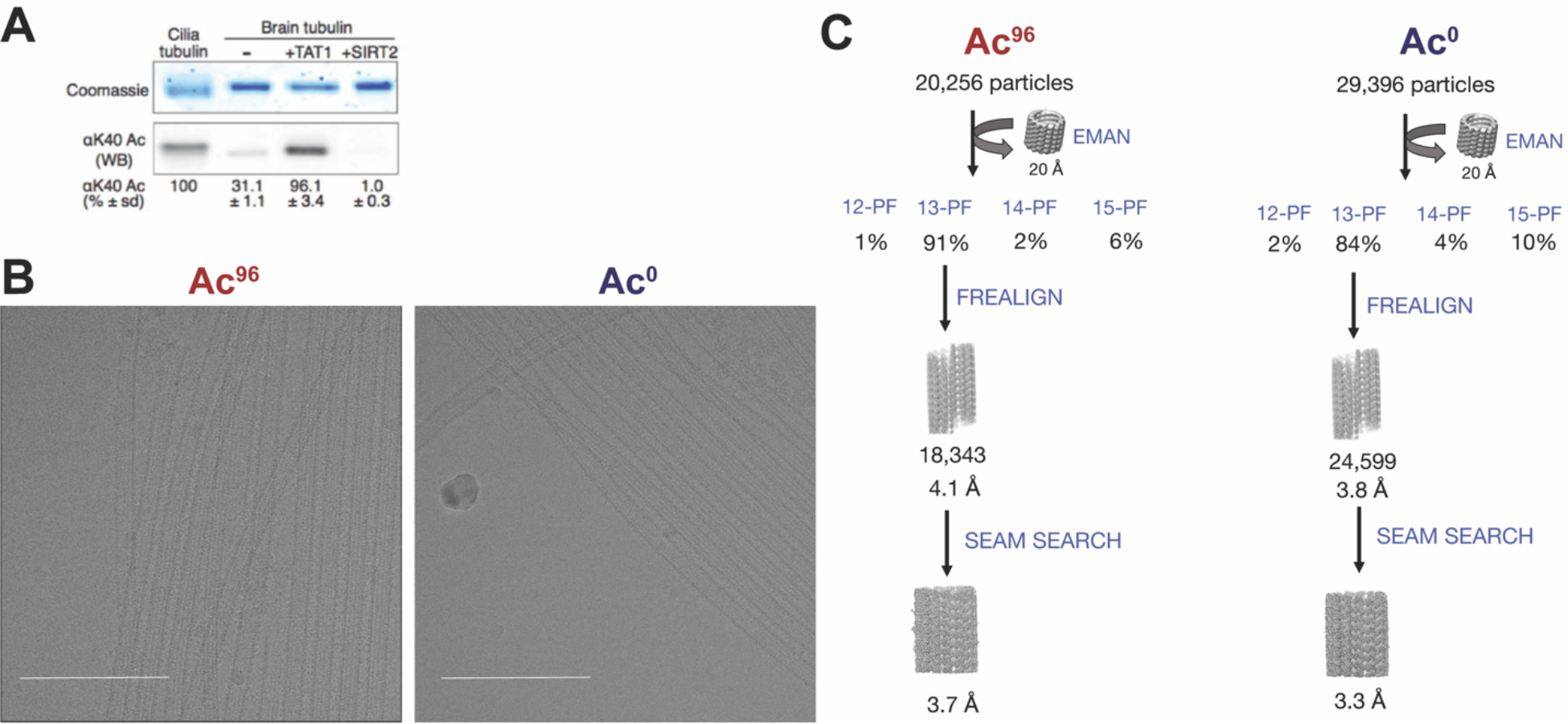
Sample preparation, data collection and image processing of acetylated and deacetylated MT samples. **(a)** Ac^96^ and Ac^0^ tubulin preparations were produced by treating purified mammalian brain tubulin (Ac^30^) with acetyltransferase αTAT1 and deacetylatase SIRT2. Samples were resolved on SDS-page and Coomassie-stained (top panel) or immunoblotted for αK40 acetylation (bottom panel). Axomenal preparations from Tetrahymena cilia provide a 100% acetylation calibrator. Adapted from Portran^21^. **(b)** Representative cryo-EM images of acetylated, in the left panel, and deacetylated MTs, in the right panel. Scale bar = 200 nm. Images were collected with the Titan Krios electron microscope (FEI, Hillsboro, OR) operated at 300kV and equipped with a K2 direct detector (Gatan, Pleasanton, CA). The micrographs were collected at a nominal magnification of 22,500x, resulting in a final pixel size of 1.07 Å per pixel and dose rate of 8 e-/pixel/s. **(c)** Schematic of data collection. Using EB3, we generated >80% homogeneous samples to push the resolution to ~3.5 Å.

**Supplemental Figure 3.**
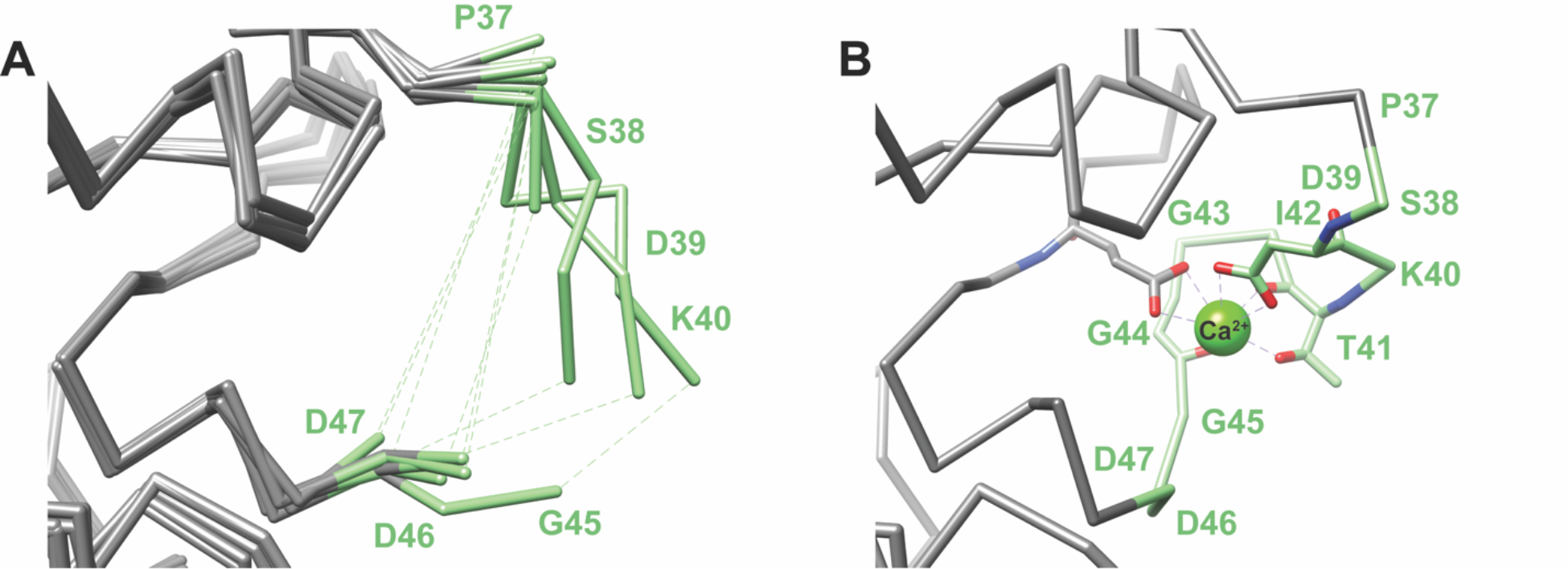
Previous proposed αK40 loop models. **(a)** Published PDBs with incomplete models of the loop: 5NQU (Chain A), 5EYP (Chain A), 3RYC (Chain A), 3RYC (Chain C), 5NQT (Chain A), 3RYI (Chain A), 3RYI (Chain A), 3RYF (Chain A), 3RYF (Chain C). **(b)** Example of the a published PDB with the complete loop stabilized by calcium: 5YL4 (Chain C).

**Supplemental Figure 4.**
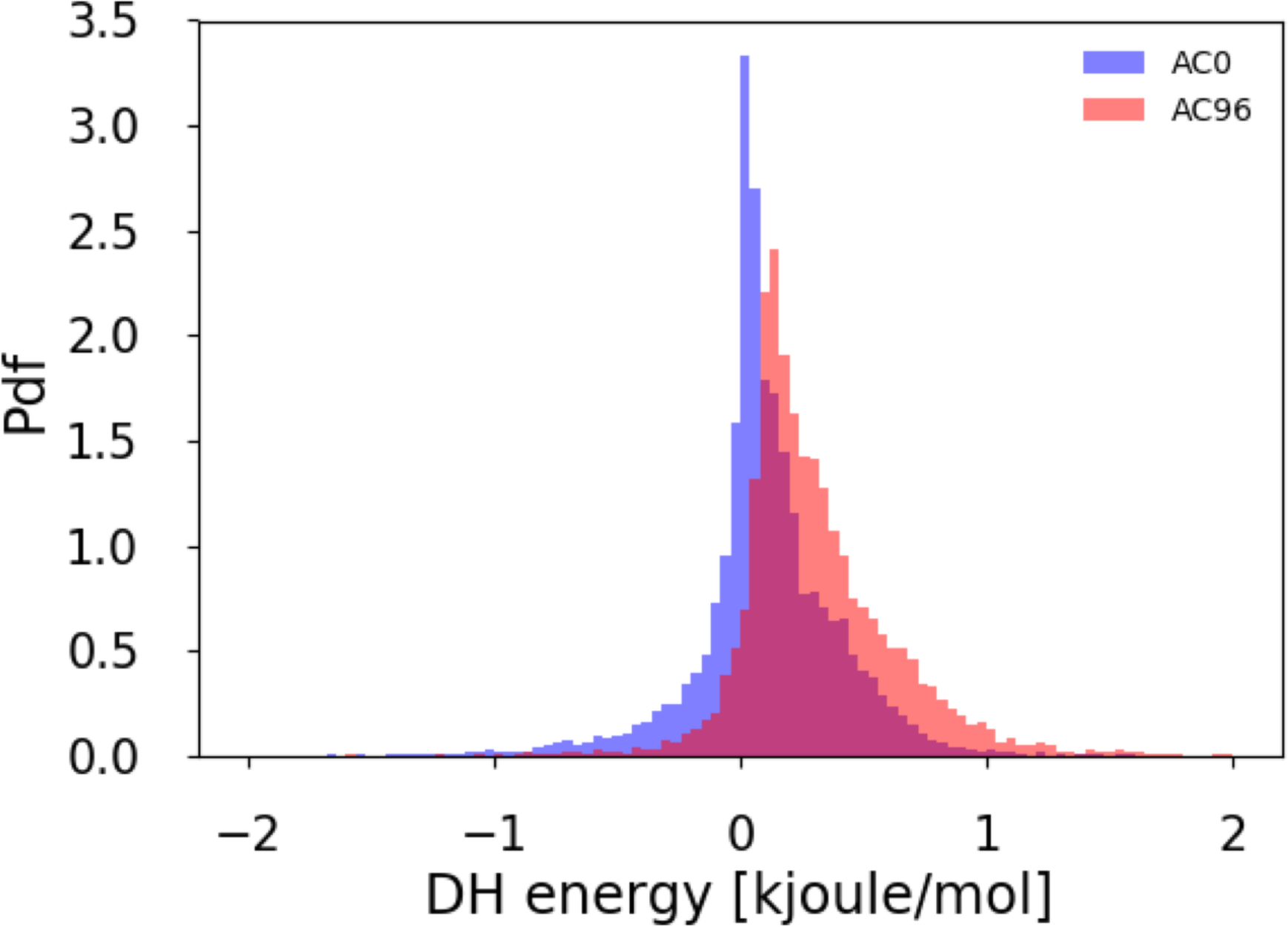
Acetylation weakens lateral interactions. By analyzing the distribution of Debye-Hückel (DH) electrostatic energy between adjacent α-subunits across the AC0 (blue) and AC96 ensembles (red), we find that acetylation weakens lateral interactions. The DH energy is calculated between the following two groups of atoms: (i) all atoms in residue range 30-60 of chain A (α1 subunit) and (ii) all atoms in residue range 200-380 of chain E (α2 subunit) in PDBs **<u>XXYA</u>** and **<u>XXYB</u>.** The plot shows the probability density function, or Pdf, as a function of the DH interaction energy.

**Supplemental Figure 5.**
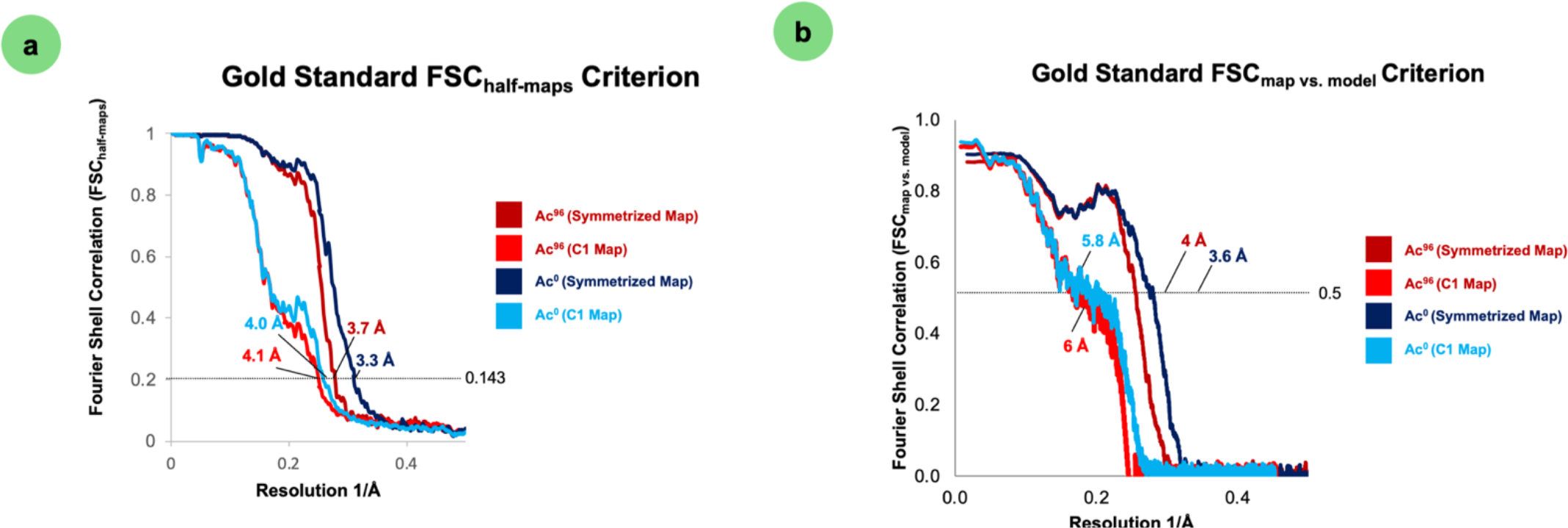
Fourier Shell Correlation Plots. The FSC_half-map_ resolution, using **0.143** as the gold standard criterion, represents how well the two half-maps from each dataset correlate as a function of spatial frequency. The two half-maps were generated by dividing the final dataset into two independent 3D-reconstructions. The FSC_map vs.model_ resolution, using **0.5** as the gold standard criterion, represents how well the final map correlated with the refined atomic model. All plots were generated in PHENIX.

**SUPPLEMENTAL TABLE 1.**
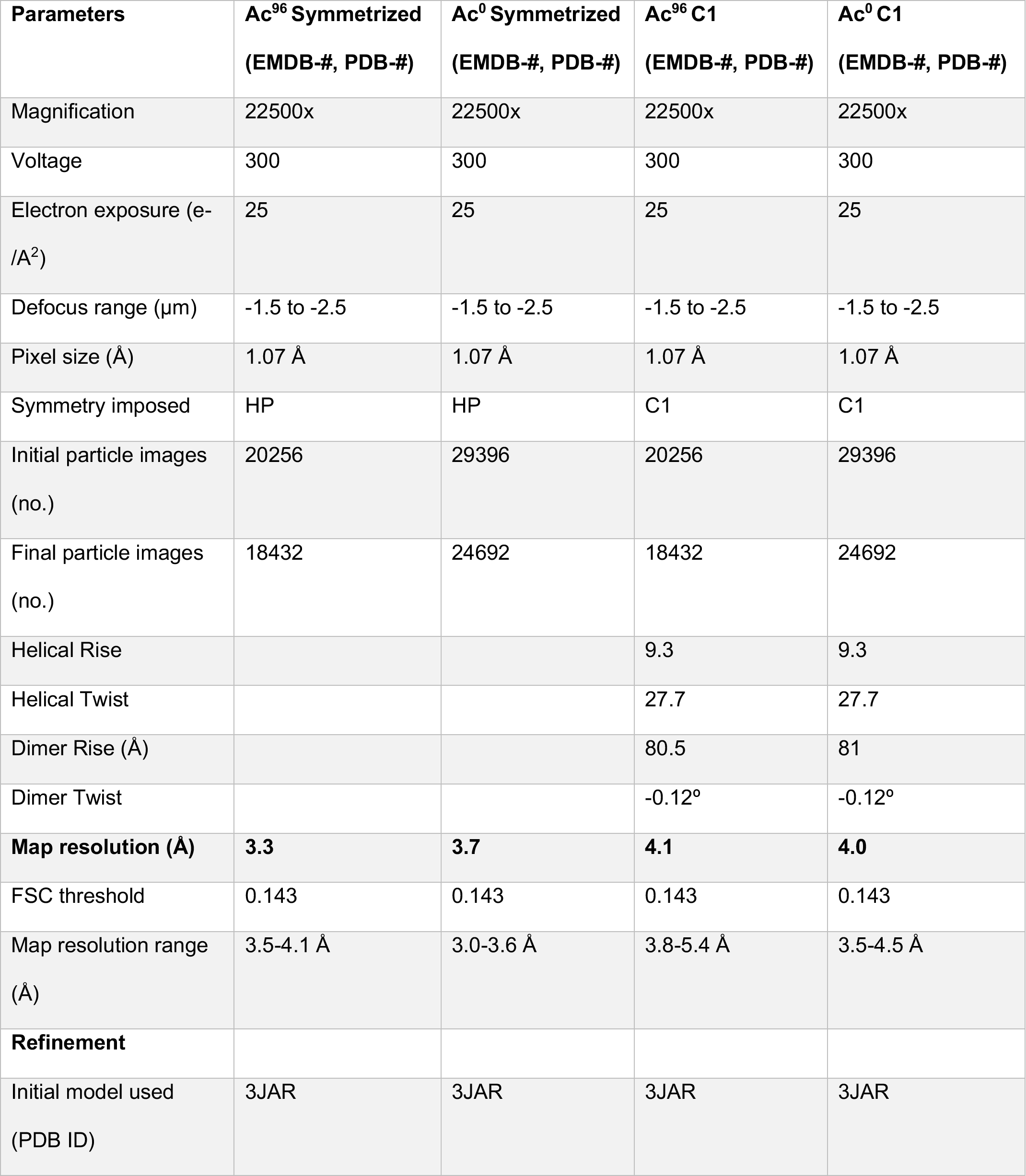
**CryoEM data collection, refinement parameters, and validation statistics**

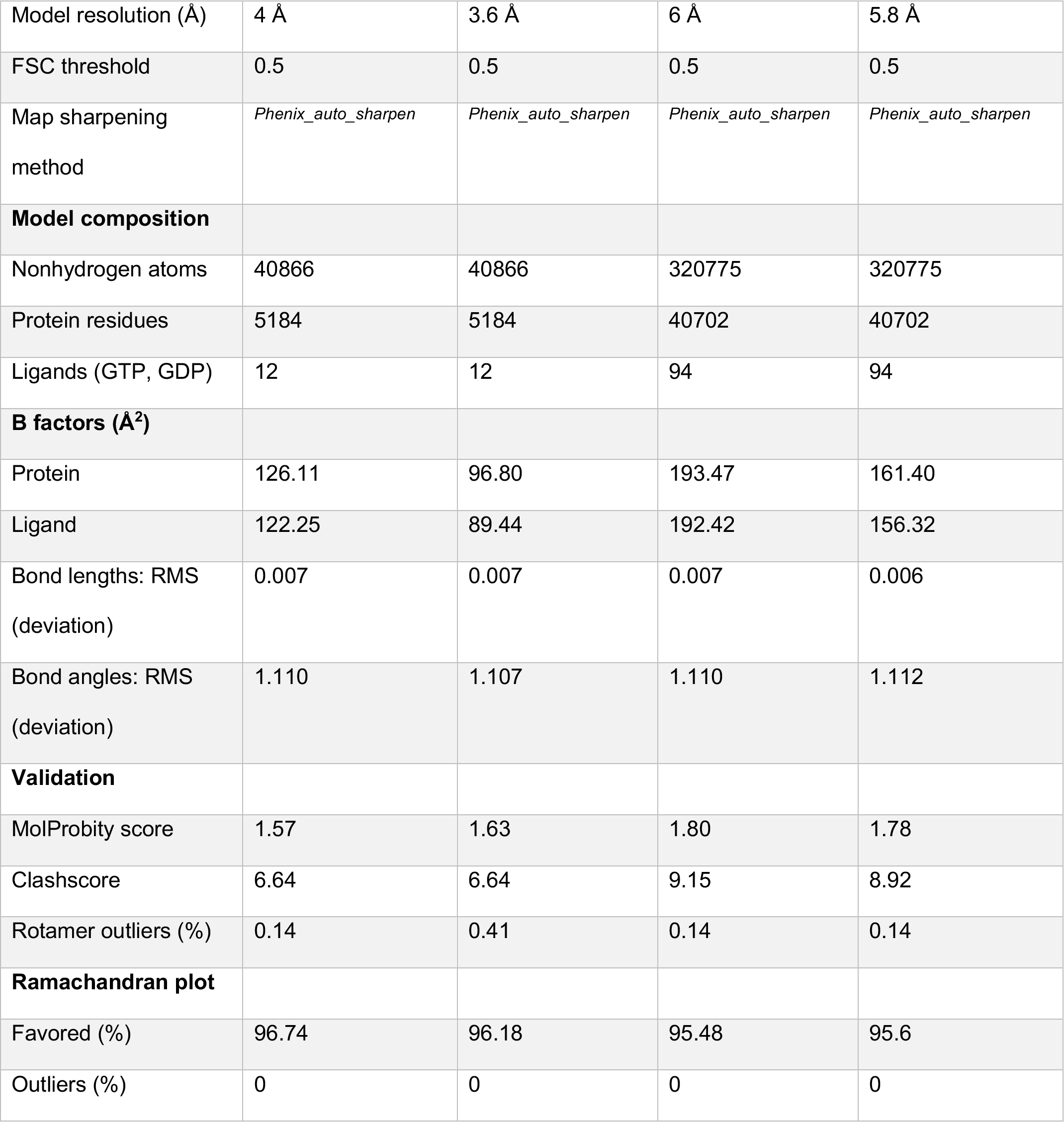

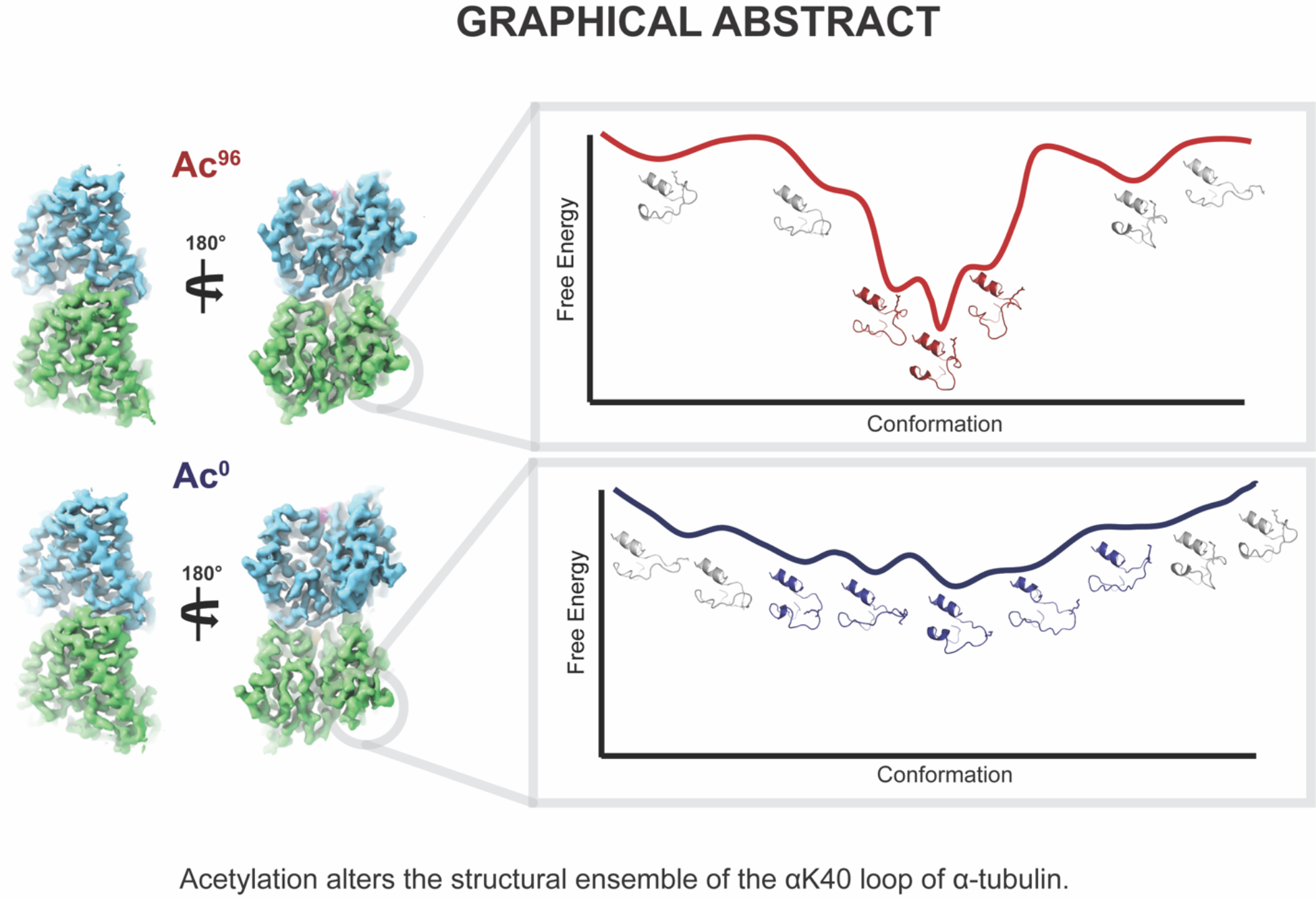

## ABBREVIATIONS

MT: microtubule
PF: protofilament
PTM: post-translational modification
MAP: microtubule-associated protein
αTAT1: acetyltransferase TAT1 for α-tubulin
SIRT2: deacetylase SIRT2.

